# Territory acquisition mediates the influence of predators and climate on juvenile red squirrel survival

**DOI:** 10.1101/594036

**Authors:** Jack G. Hendrix, David N. Fisher, April Robin Martinig, Stan Boutin, Ben Dantzer, Jeffrey E. Lane, Andrew G. McAdam

## Abstract

1) Juvenile survival to first breeding is a key life history stage. Survival through this period can be particularly challenging when it coincides with harsh environmental conditions like winter climate or food scarcity, and so cohort survival can be highly variable. However, the small size and dispersive nature of juveniles makes studying their survival difficult.
2) In territorial species, a key life history event is the acquisition of a territory. A territory is expected to enhance survival, but how it does so, possibly through mediating mortality, is not often identified. We tested how the timing of territory acquisition influenced the survival of juvenile North American red squirrels *Tamiasciurus hudsonicus*, hereafter red squirrels, and how the timing of this event mediated sources of mortality. We hypothesized that securing a territory prior to the caching season would reduce juvenile susceptibility to predation or climatic factors over winter.
3) Using 27 years of data on the survival of individually-marked juvenile red squirrels, we tested how the timing of territory acquisition influenced survival, whether the population density of red squirrel predators and mean temperature over winter were related to individual survival probability, and if territory ownership mediated these effects.
4) Juvenile survival was lower in years of high predator abundance and in colder winters. Autumn territory owners were less susceptible to lynx *Lynx canadensis*, and possibly mustelid *Mustela* and *Martes* spp., predation. Autumn territory owners had lower survival in colder winters, while non-owners had higher survival in cold winters.
5) Our results show how the timing of a life history event like territory acquisition can directly affect survival and also mediate the effects of biotic and abiotic factors later in life. This engenders a better understanding of the fitness consequences of the timing of key life history events.

## Introduction

Survival as a juvenile, when individuals are no longer completely dependent on the parent but not yet sexually mature, is a crucial life history stage for all taxa (Ferguson & Fox, 1984; Gaillard, Festa-Bianchet, & Yoccoz, 1998; Searcy & Sponaugle, 2001). A large component of reproductive success is surviving to sexual maturity, hence juvenile survival can be a key determinant of lifetime fitness, and so variation in survival can dictate population dynamics (McAdam, Boutin, Sykes, & Humphries, 2007; Oli & Dobson, 2003). For example, rates of juvenile survival in large herbivores are highly variable year-to-year, and despite the fact that they do not determine population growth rates, they may be the key determinant of population dynamics (Gaillard, Festa-Bianchet, Yoccoz, Loison, & Toïgo, 2000). Understanding the causes of variation in juvenile survival and the selection this facilitates therefore shapes how we expect populations to change over time.

The time period between juvenile independence and first breeding poses particular challenges to survival as juvenile mortality is often high during this period. Climatic factors can have strong effects on survival of juveniles (Fuller, Stebbins, & Dyke, 1969; Schorr, Lukacs, & Florant, 2009) through a combination of limited food availability and increased thermoregulatory costs (Jackson, Trayhurn, & Speakman, 2001; Rödel et al., 2004), particularly over winter. Due to their small size and lack of experience, as well as their dispersive nature, juveniles can also be particularly vulnerable to predation (Garrett & Franklin, 1988; Rödel et al., 2015). Various behavioural and physiological responses such as adjusting metabolic rate (Wunder, Dobkin, & Gettinger, 1977), reducing activity (Merritt, 1986), or food caching (Morrison, Pelchat, Donahue, & Hik, 2009) can mitigate this risk. Understanding how these mediating traits alter juvenile survival is necessary to understand how selection shapes phenotypes.

The acquisition of a territory is a key life history event that can mediate sources of mortality in some species, by providing access to space, refuges and food stores. Timing of life history stages, such as birth or hatching (Rodríguez, van Noordwijk, Álvarez, & Barba, 2016), or developmental rate (van der Jeugd & Larsson, 1998) can have strong effects on survival at later life stages (O’Connor, Norris, Crossin, & Cooke, 2014). Territory acquisition is one such event: predation risk is elevated while searching for territories (Larsen & Boutin, 1994), and territory ownership also leads to increased food availability, particularly in food caching species. Earlier acquisition of a territory can, therefore, improve the probability of survival by reducing these risks sooner in life. It is well known that acquiring a territory provides benefits (reviewed in: Carpenter, 1987; e.g. Whitham, 1986). However, despite the potential importance of understanding how the timing of territory acquisition modifies juvenile survival and mediates sources of mortality, documenting the consequences for mortality risks of the timing of territory acquisition has not occurred, in part due to the difficulty in collecting such data.

North American red squirrels *Tamiasciurus hudsonicus* are an ideal organism to study how the timing of territory acquisition influences survival and mediates sources of juvenile mortality. Red squirrels in Yukon, Canada defend exclusive individual territories with a central cache of white spruce *Picea glauca* cones, their primary food source (Boutin & Schweiger, 1988; Fletcher et al., 2013). Holding a territory with a cache of food is considered necessary for red squirrels in these populations to survive over winter (Larsen & Boutin, 1994; Smith, 1968), as cached resources are essential for annual survival and reproduction (Fletcher et al., 2013; LaMontagne et al., 2013). White spruce cones ripen in early August and caching begins shortly thereafter and finishes at the end of September (Fletcher et al., 2010). Those juveniles with territories before this ripening begins are able to take advantage of that year’s cone crop and increase their hoard size, whereas those that settle on territories later in the season can only take over what is left from the previous owner (Fisher et al., 2019) and have no opportunity to secure further resources before winter.

Juvenile annual over winter mortality is high, with an average of 73.6 % not surviving their first winter (McAdam et al., 2007), but this is highly variable annually (57 - 97 %) (McAdam & Boutin, 2003). Annual adult mortality in this population is low (20 % for two year old females; this steadily increases with age), thus much of the variation in lifetime reproductive success is linked to juvenile over winter mortality (McAdam et al., 2007). Acquiring a territory is therefore a key life history event. However, the main causes of juvenile mortality, and how they are influenced by the timing of territory acquisition, are not known.

Observational studies, while relatively limited, have identified lynx *Lynx canadensis* (Stuart-Smith & Boutin, 1995), goshawks *Accipiter gentilis* (Larsen & Boutin, 1994), and mustelids (Kerr & Descamps, 2008; O’Donoghue, Boutin, Hofer, & Boonstra, 2001) as predators of juvenile red squirrels (Goheen & Swihart, 2005; Haines et al., 2018; Smith, 1968; Steele, 1998). Owning a territory, and thus having access to nests or tunnels, could act as spatial refugia and reduce vulnerability to predators (Cowlishaw, 1997; Everett & Ruiz, 1993). Furthermore, red squirrels with smaller caches have lower over winter survival (LaMontagne et al., 2013; Larivée, Boutin, Speakman, McAdam, & Humphries, 2010), suggesting that resource limitation is a source of over winter mortality. Owning a territory, and so regular use of nests, would provide thermal refugia during low temperatures (Greenwood & Harvey, 1982; Studd, Boutin, McAdam, Krebs, & Humphries, 2015). It therefore seems that a territory could both directly influence survival and change the suite of selection pressures that act on a juvenile red squirrel.

We aimed to better understand how territory acquisition affects juvenile over winter survival and mediates sources of mortality. To do so we used 27 years of longitudinal data to assess how holding a territory before autumn influences survival and the susceptibility of a juvenile to predation or low temperatures over winter.

Our first hypothesis was that earlier territory acquisition would give higher over winter survival compared to later territory acquisition (Berteaux & Boutin, 2000). We further hypothesized that cold temperatures and predators pose a mortality risk, so that over winter survival of juveniles would be lower in colder winters and when predators are abundant. Our key hypothesis is that timing of territory acquisition would moderate these effects, so that juveniles obtaining territories before autumn are less susceptible to predators (e.g. Cowlishaw 1997) and adverse weather (e.g. Greenwood and Harvey 1982) over winter.

## Materials and Methods

### Data collection

Our study was part of the Kluane Red Squirrel Project, an ongoing long-term study of a wild population of North American red squirrels within Champagne and Aishihik First Nations traditional territory along the Alaska Highway in southwestern Yukon, Canada (61° N, 138° W). We collected data from two study areas (~ 40 hectares each) separated by the Alaska Highway from 1989 to 2015. We conducted population censuses biannually in May (spring) and August (autumn) to identify all individuals and assign territory ownership. We assigned territory ownership based on territorial vocalisations ("rattling"; Lair, 1990) and behavioural observations. We also identified living individuals that did not own territories in autumn through trapping and behavioural observations. Adult red squirrels rarely relocate, other than through bequeathals by mothers where all or a part of her territory is given to offspring (Berteaux & Boutin, 2000; around 19 % of females do this each year; Lane et al., 2015; Larsen & Boutin, 1994a). Average juvenile dispersal distance is short (mean = 92 to 102 m; Berteaux & Boutin, 2000; Cooper et al., 2017) relative to the size of our study areas. Juveniles born on the edge of our study areas do not have lower apparent survival than those born in the core, suggesting that dispersal outside our study areas does not bias our mortality estimates (T. D. Kerr, Boutin, LaMontagne, McAdam, & Humphries, 2007; McAdam et al., 2007).

We used Tomahawk traps (Tomahawk Live Trap Co., Tomahawk, Wisconsin, U.S.A.) baited with peanut butter on or near each individual’s midden to trap them. When handled for the first time, each individual was given numbered ear tags (Monel #1; 5 digits) with a unique combination of coloured wires or pipe cleaners to facilitate future identification without handling. We recorded body mass, sex, and reproductive status at each capture. We radio-collared reproductive females (model PD-2C, 4 g, Holohil Systems Limited, Carp, Ontario, Canada) to find nests. Females typically give birth to three pups (range: one – seven; Humphries & Boutin, 2000) in the spring (median birth date: 23 April). We removed juveniles from the nest after birth, and a second time at ~25 days old, to record litter size, pup mass, and sex, and to tag them. Growth rate (g/day) was calculated as the linear increase in mass between the nest entries. We calculated growth rates only for juveniles that weighed less than 50 g at the first nest entry and less than 100 g at the second nest entry (to ensure approximate linearity of the growth curve; McAdam & Boutin, 2003), and only for juveniles where the two weight measures were >5 days apart. Juveniles emerge from the nest around 42-50 days old and wean around 70 days (Larsen & Boutin, 1994). We considered juveniles surviving to the spring following the year of their birth to have recruited into the population. This research was approved by the University of Guelph Animal Care Committee (AUP 1807), the University of Alberta Animal Care and Use Committee for Biosciences and the University of Michigan Institutional Animal Care and Use Committee (PRO00007805).

### Predator and temperature data collection

Our available temperature and predator data are annual, regional measures, so for this analysis we treated all juveniles born in the same year as experiencing the same conditions. We obtained monthly temperature records from Environment Canada’s online historical weather database for the Haines Junction weather station (Climate ID 2100630, 60.77° N, 137.57° W), approximately 35 km SE of our study area. We used mean temperature over winter, as we expected that climate would primarily influence over winter survival by increasing thermoregulatory costs as opposed to extreme weather events or precipitation. We averaged the monthly temperatures from October of a juvenile’s birth year to the following March to obtain an annual average winter temperature.

We considered potential mammalian predators: mustelids (short-tailed weasel *Mustela erminea*, least weasel *M. nivalis*, and marten *Martes americana*) and lynx *Lynx canadensis*. We obtained abundance data for predators and their alternate prey from population monitoring in our region, first as part of the Kluane Boreal Forest Ecosystem Project (Krebs, 2001), and after 1996 as part of the Community Ecological Monitoring Program (available at http://www.zoology.ubc.ca/~krebs/kluane.html). Repeated track counts along set transects over winter provided an estimate of species abundance as the mean number of tracks per 100 km of transect. We used the sum of short-tailed weasel, least weasel, and marten tracks as the total mustelid abundance for each year. The population densities of snowshoe hares *Lepus americanus* and red-backed voles *Myodes rutilis* were estimated with live trapping and mark-recapture, providing measures of alternate prey availability for these predators. These combinations were chosen as lynx are known hare specialists (O’Donoghue, Boutin, Krebs, Murray, & Hofer, 1998), while weasels (the majority of the mustelids) are known vole specialists (Boonstra & Krebs, 2006), and both populations follow the cycles of their preferred prey (Boutin et al. 1995). While birds of prey such as goshawks *A. gentilis* are known predators of red squirrels (Larsen & Boutin, 1994), we were not able to include them in our analysis as population counts over time are not available. Such birds of prey primarily prey on snowshoe hares, and so their population sizes typically tack those of the hares, as lynx population sizes do (Boutin et al. 1995). Therefore, the effect of lynx abundance may somewhat represent the overall effect of snowshoe hare predators on red squirrels.

### Statistical analyses of survival

We used a binomial mixed effects model to test how predation and temperature interacted with autumn territory ownership to affect juvenile survival over winter. From 1989 to 2015, our analysis considered whether those juveniles that survived to the beginning of August (n = 1305 squirrels) were still alive the following spring.

We included several factors previously shown to affect juvenile survival in our system (Descamps, Boutin, Berteaux, & Gaillard, 2008): these included squirrel population density (number of adults within a set 38 ha area; this area was consistent over the entire study period), spruce cone availability (annual index of cones produced on a consistent subset of trees on each study area; see: LaMontagne et al., 2005), a fixed effect of study area to account for any differences between the two study areas, birth date, growth rate, and sex. Growth rate, birth date, adult population density, and cone availability were standardized as z-scores for each study area in each year. This improves model convergence and interpretability of regression coefficients (Schielzeth, 2010).

For our main question – does territory ownership mediate how predators and climate affect survival – we included territory ownership in autumn as a binary predictor with temperature and predator abundance as numeric predictors, and fit interactions between autumn ownership and each of temperature, lynx, and mustelid abundances separately. We included separate interactions between the abundance of lynx and snowshoe hares, and mustelids and voles, so the effect of predators on red squirrels depended on the availability of preferred prey. Temperature and species abundances were standardized as z-scores across years. Finally, we included random effects of litter identity and year to account for variation in survival due to sibling and maternal interactions, as well as otherwise unaccounted for annual variation.

We also fitted a separate model with interactions of juvenile birth date and growth rate with predators and temperature, to determine whether these traits influence these sources of mortality. We present these results in the supporting information (Table S1): we found no evidence of predator abundance or temperature over winter acting as agents of selection on either of these traits.

We conducted all statistical analyses using R version 3.3.3 (R Core Team 2017), with the packages lme4 (version 1.1-19; Bates et al., 2015), and lmerTest (version 2.0-33; Kuznetsova, Brockhoff, & Christensen 2016). Reported estimates are means ± SE.

## Results

### Over winter survival

Among juveniles alive in August between 1989 - 2015, an average of 60 % survived to the following spring, but this was highly variable annually (21.4 - 94.1 %; Table 1). Juvenile survival was higher with increased cone availability (β = 0.38 ± 0.11, *z* = 3.45, *P* < 0.001; Table 2) and years of lower adult population density (β = −0.69 ± 0.15, *z* = −4.45, *P* < 0.001). Female juveniles were more likely to survive over winter than males (β = 0.49 ± 0.16, *z* = 3.1, *P* = 0.002), as were juveniles with higher growth rates (β = 0.22 ± 0.10, *z* = 2.13, *P* = 0.033). Birth date had no effect on over winter survival (β = −0.01 ± 0.09, *z* = −0.08, *P* = 0.936), nor were there any differences between study areas (β = 0.19 ± 0.18, *z* = 1.06, *P* = 0.289). The random effect of litter ID explained a significant amount of variation (σ^2^ = 0.665; likelihood ratio test X^2^ = 7.867, *df* = 20, *P* = 0.005), but the random year effect did not contribute significantly to the model.

**Table 1.**
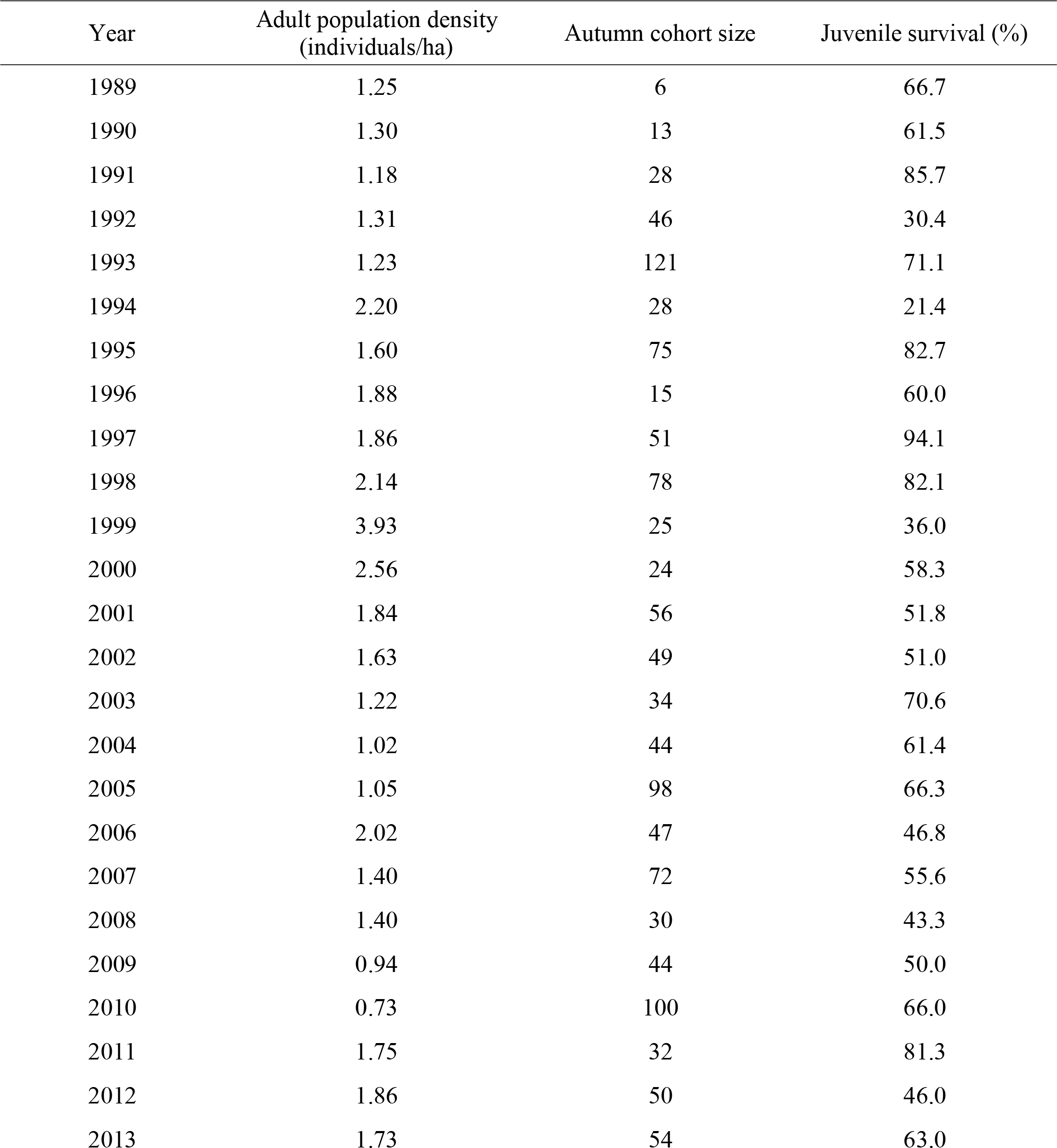
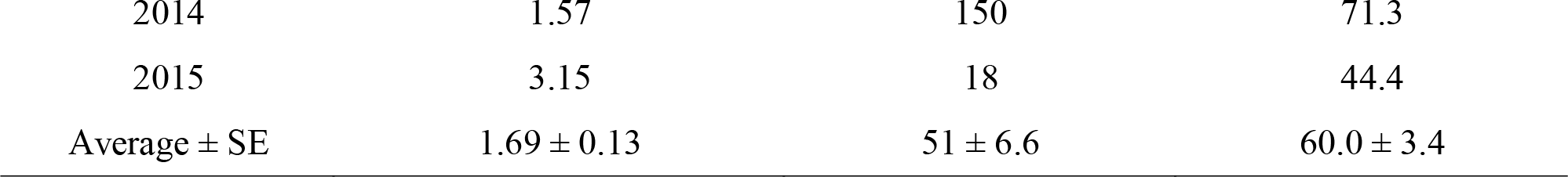
Probability of over winter survival for juvenile red squirrels alive in August 1989 – 2015 (*n* = 1305), with adult density for each year (individuals/ha) and number of juveniles alive in autumn (cohort size).

| Year | Adult population density<br>(individuals/ha) | Autumn cohort size | Juvenile survival (%) |
| --- | --- | --- | --- |
| 1989 | 1.25 | 6 | 66.7 |
| 1990 | 1.30 | 13 | 61.5 |
| 1991 | 1.18 | 28 | 85.7 |
| 1992 | 1.31 | 46 | 30.4 |
| 1993 | 1.23 | 121 | 71.1 |
| 1994 | 2.20 | 28 | 21.4 |
| 1995 | 1.60 | 75 | 82.7 |
| 1996 | 1.88 | 15 | 60.0 |
| 1997 | 1.86 | 51 | 94.1 |
| 1998 | 2.14 | 78 | 82.1 |
| 1999 | 3.93 | 25 | 36.0 |
| 2000 | 2.56 | 24 | 58.3 |
| 2001 | 1.84 | 56 | 51.8 |
| 2002 | 1.63 | 49 | 51.0 |
| 2003 | 1.22 | 34 | 70.6 |
| 2004 | 1.02 | 44 | 61.4 |
| 2005 | 1.05 | 98 | 66.3 |
| 2006 | 2.02 | 47 | 46.8 |
| 2007 | 1.40 | 72 | 55.6 |
| 2008 | 1.40 | 30 | 43.3 |
| 2009 | 0.94 | 44 | 50.0 |
| 2010 | 0.73 | 100 | 66.0 |
| 2011 | 1.75 | 32 | 81.3 |
| 2012 | 1.86 | 50 | 46.0 |
| 2013 | 1.73 | 54 | 63.0 |
| 2014 | 1.57 | 150 | 71.3 |
| 2015 | 3.15 | 18 | 44.4 |
| Average ± SE | $1.69 \pm 0.13$ | $51 \pm 6.6$ | $60.0 \pm 3.4$ |

**Table 2.** Mixed effects binomial model of juveniles red squirrel over winter survival (*n* = 1305), testing whether territory ownership by autumn mediates effects of predators and temperature on over winter survival, including random effects of litter ID and year (conditional R^2^ = 0.44).

| Term | Estimate ± SE | $z$ | $P$ |
| --- | --- | --- | --- |
| Std. density | $-0.69 \pm 0.15$ | -4.45 | < 0.001 |
| Std. cones | $0.38 \pm 0.11$ | 3.45 | < 0.001 |
| Std. growth rate | $0.22 \pm 0.10$ | 2.13 | 0.033 |
| Std. birth date | $-0.01 \pm 0.09$ | -0.08 | 0.936 |
| Grid (SU) | $0.19 \pm 0.18$ | 1.06 | 0.289 |
| Sex (male) | $-0.49 \pm 0.16$ | -3.1 | 0.002 |
| Autumn owner (yes) | $2.78 \pm 0.23$ | 12.06 | < 0.001 |
| Std. lynx | $-0.68 \pm 0.28$ | -2.41 | 0.016 |
| Std. hares | $0.40 \pm 0.20$ | 1.99 | 0.046 |
| Std. mustelid | $-0.38 \pm 0.14$ | -2.7 | 0.007 |
| Std. voles | $-0.59 \pm 0.13$ | -4.57 | < 0.001 |
| Std. temperature | $-0.35 \pm 0.18$ | -1.99 | 0.047 |
| Std. lynx : Std. hares | $0.12 \pm 0.09$ | 1.32 | 0.187 |
| Std. mustelid : Std. voles | $0.14 \pm 0.11$ | 1.27 | 0.203 |
| Autumn owner (yes) : Std. lynx | $0.99 \pm 0.23$ | 4.22 | < 0.001 |
| Autumn owner (yes) : Std. mustelid | $0.31 \pm 0.18$ | 1.75 | 0.080 |
| Autumn owner (yes) : Std. temperature | $1.11 \pm 0.21$ | 5.31 | < 0.001 |
| Random effects | Variance |  |  |
| Litter ID | 0.665 |  |  |
| Year | 0.000 |  |  |

### Territory ownership and over winter survival

Sixty-one percent of juveniles alive in August owned a territory, and these juveniles were more likely (79 %) to survive over winter than those who did not (33 %; β = 2.78 ± 0.23, *z* = 12.06, *P* < 0.001). Juveniles without territories in August were less likely to survive in years of high lynx (β = −0.68 ± 0.28, *z* = −2.41, *P* = 0.016) and mustelid (β = −0.38 ± 0.14, *z* = −2.70, *P* = 0.007) abundance. There was a significant interaction between lynx abundance and territory ownership (β = 0.99 ± 0.23, *z* = 4.22, *P* < 0.001; Fig 1); increased lynx abundance had no effect on the over winter survival of juveniles that held territories by autumn. The mustelid – owner interaction was not significant, but in the same direction as for lynx, with mustelid abundance having a weaker effect on territory owners (β = 0.31 ± 0.18, *z* = 1.75, *P* = 0.080; Fig 2). The effects of predators on juvenile survival did not depend on the abundance of alternate prey (lynx x hare *P* = 0.187; mustelid x vole *P* = 0.203), although both the hare (β = 0.40 ± 0.20, *z* = 1.99, *P* = 0.046) and vole (β = −0.59 ± 0.13, *z* = −4.57, *P* < 0.001) main effects were significant.

**Figure 1.**
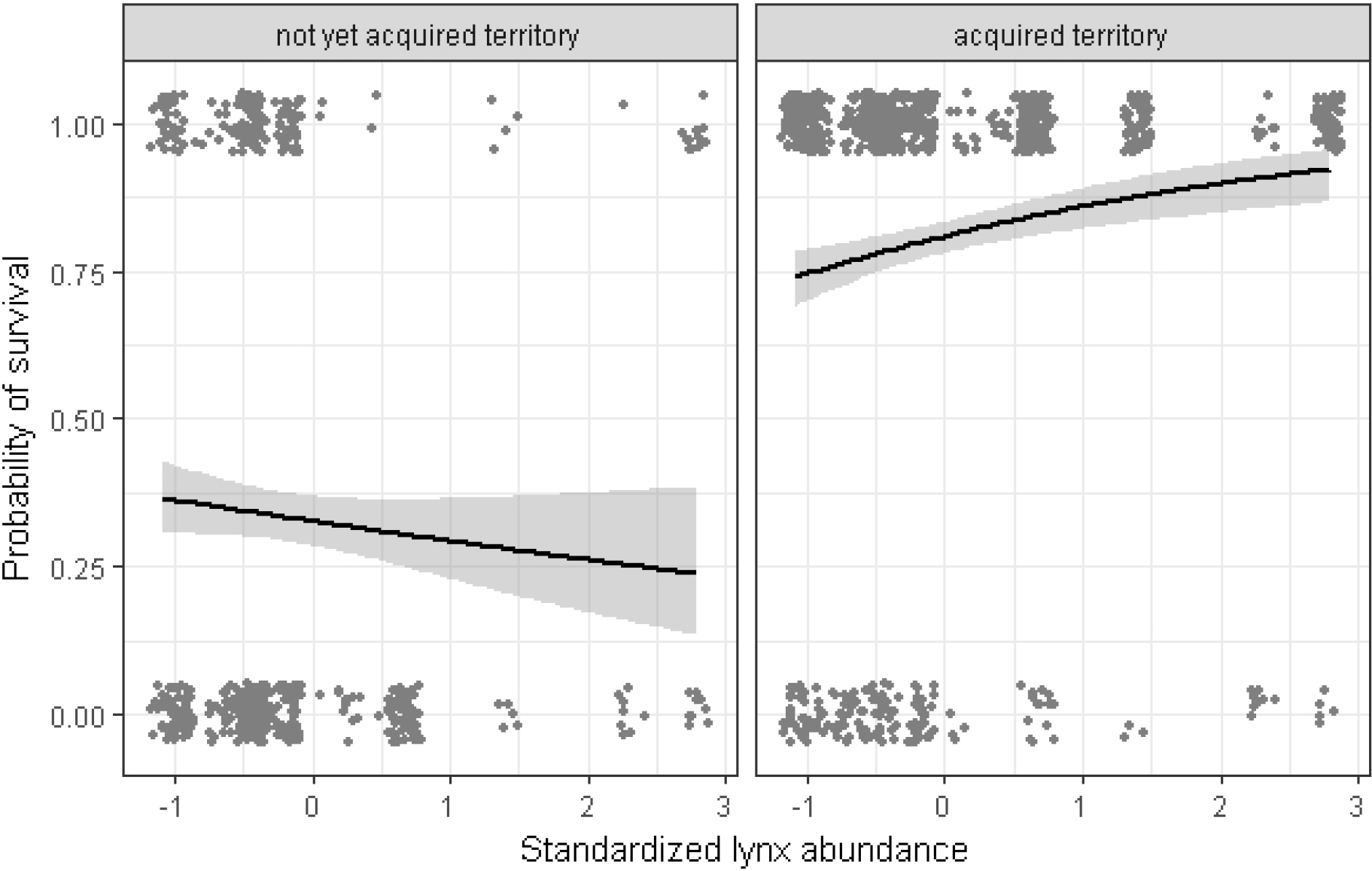
Over winter survival of juveniles (*n* = 1305) that had or had not acquired a territory by autumn. Juveniles without territories had lower survival when lynx were abundant (non-owners: β = −0.68 ± 0.28, *z* = −2.41, *P* = 0.016), whereas the survival of juveniles with territories was unaffected by lynx abundance (owners: β = 0.31 ± 0.21, *z* = 1.49, *P* = 0.14; interaction β: = 0.99 ± 0.23, *z* = 4.22, *P* < 0.001).

**Figure 2.**
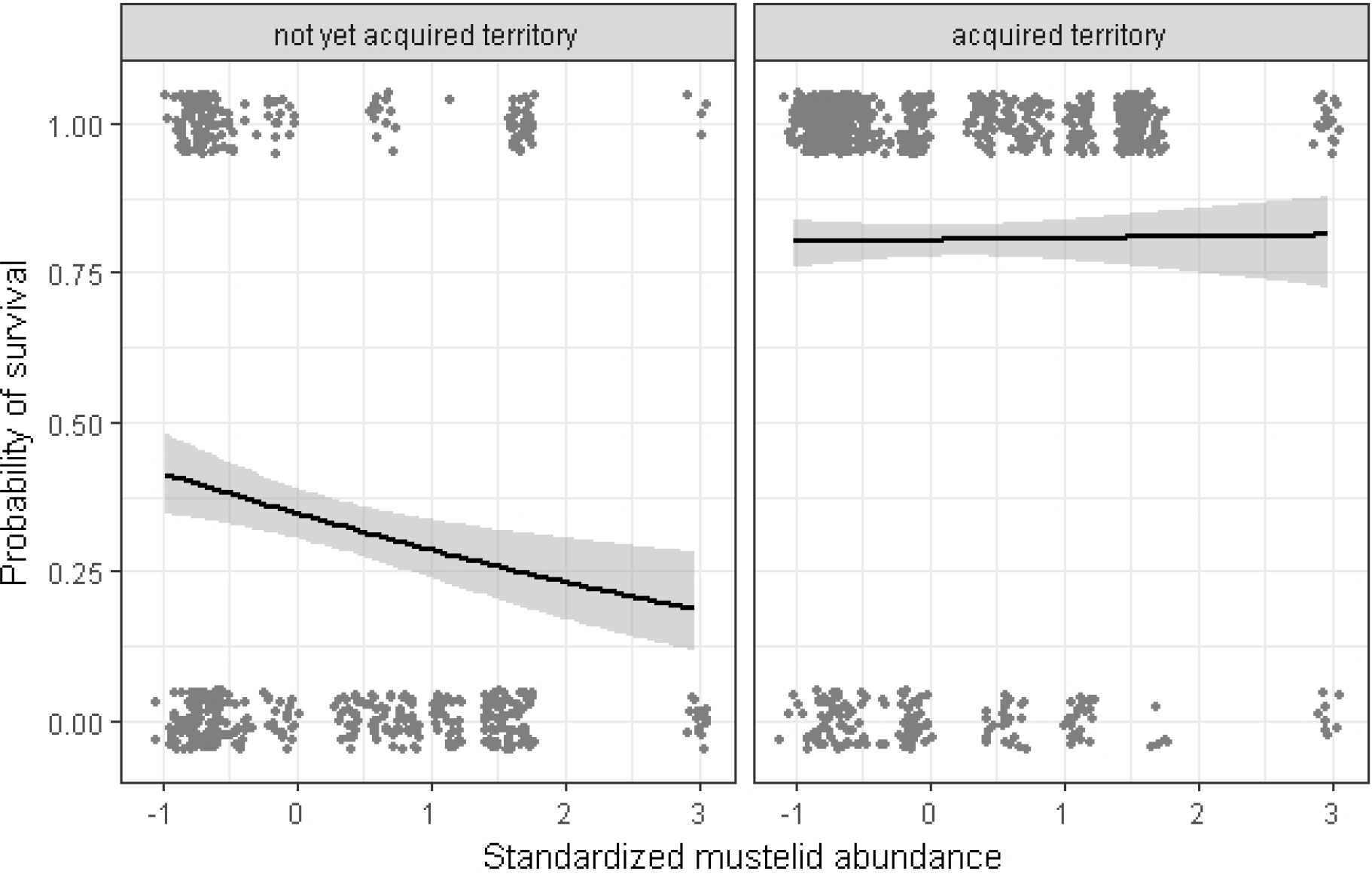
Over winter survival of juveniles (*n* = 1305) that had or had not acquired a territory by autumn was lower when mustelids were abundant. Juveniles without territories by autumn were somewhat more affected by mustelid abundance (non-owners: β = −0.38 ± 0.14, *z* = −2.70, *P* = 0.007) than territory owners (owners: β = −0.07 ± 0.14, *z* = −0.49, *P* = 0.624; interaction β = 0.31 ± 0.18, *z* = 1.75, *P* = 0.080).

Temperature had opposing effects on survival for juveniles with and without territories by autumn (Fig 3). Juveniles without territories by autumn were less likely to survive warm winters (β = −0.35 ± 0.18, *z* = −1.99, *P* = 0.047), but this effect reversed for autumn territory owners (interaction β = 1.11 ± 0.21, *z* = 5.31, *P* < 0.001), which were more likely to survive warm winters (β = 0.76 ± 0.13, *z* = 5.31, *P* < 0.001).

**Figure 3.**
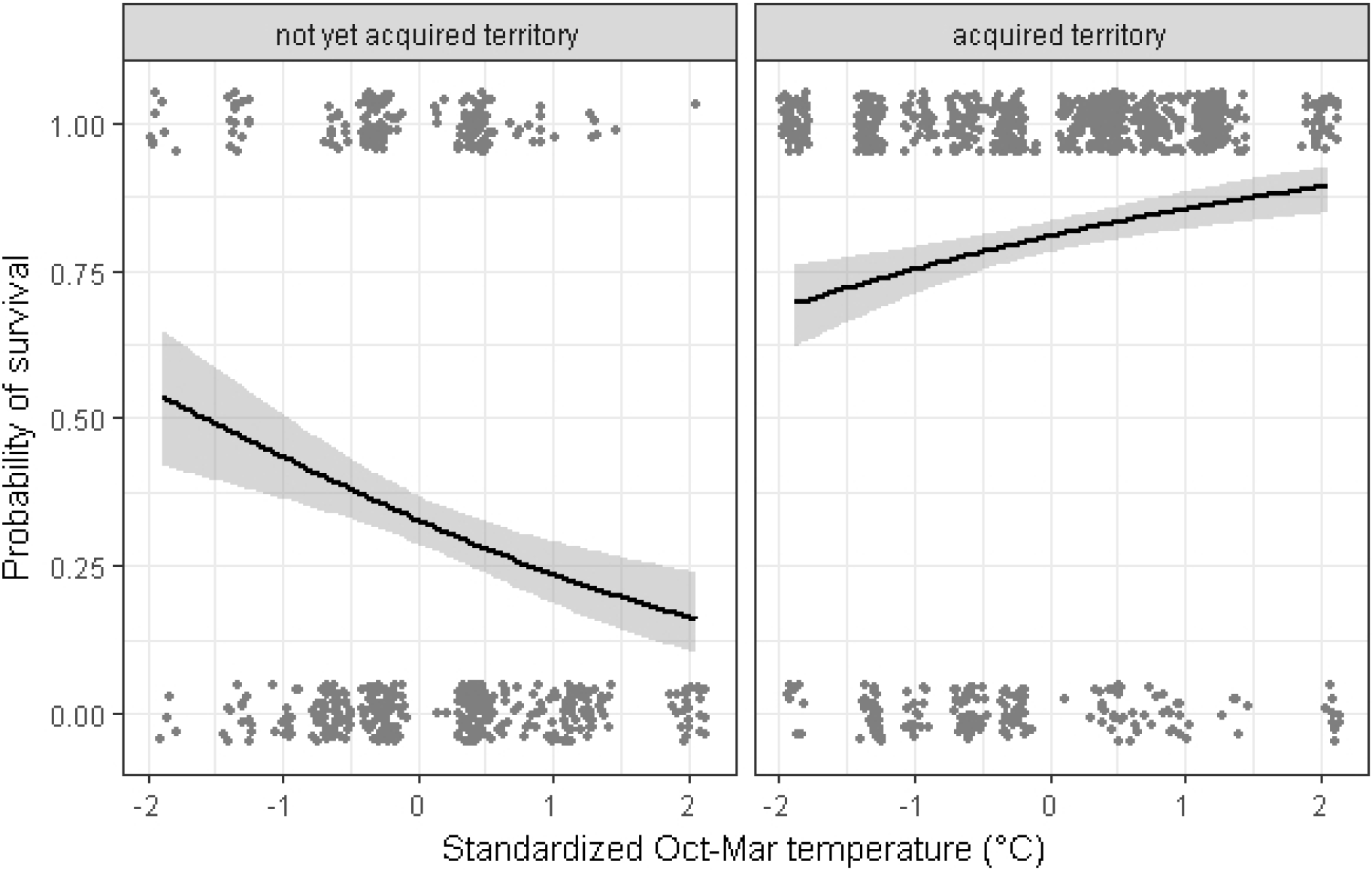
Over winter survival of juveniles (*n* = 1305) that had or had not acquired a territory by autumn. Autumn territory owners survived better in warmer years (owners: β = 0.76 ± 0.13, *z* = 5.87, *P* < 0.001), whereas warmer winters increased mortality of juveniles without territories at this time (non-owners: β = −0.35 ± 0.18, *z* = −1.99, *P* = 0.047; interaction β: = 1.11 ± 0.21, *z* = 5.31, *P* < 0.001).

## Discussion

Juveniles that acquired territories earlier in the year were far more likely to survive the winter than those that had not yet found a territory before autumn. Average survival of juveniles that acquired territories before the start of cone caching (79 %) was comparable to survival of early-life adults in this population (80 %; McAdam et al., 2007). Juveniles without territories by autumn had much lower survival (33 %), as they may have never acquired a territory and so perished or acquired one late but lost the opportunity to cache resources in it, and so did not have a large enough stockpile of resources to survive over winter. Their lower estimated survival is unlikely to be because the juveniles without a territory had in fact moved off our study area, as survival is equal between juveniles originating from the centre of the study area and those at the edge (T. D. Kerr et al., 2007).

Territory ownership also affected how susceptible juveniles were to predators and weather over winter. Juveniles without territories by autumn were more susceptible to predators than those that had already settled. Territory ownership provides access to arboreal nests, midden tunnels, and increased familiarity with the local habitat (Clarke et al., 1993). Juveniles without territories by autumn may be travelling more through potentially high-risk environments as they forage for food or search for territories over winter, thereby increasing their vulnerability to predators (Garrett & Franklin, 1988; Metzgar, 1967). Higher rates of litter loss in red squirrels during years of high mustelid abundance (Studd et al., 2015) suggests that mustelids can access red squirrel nests (and likely tunnels) whereas lynx may be more effectively deterred by these structures. This may explain why the relationship between mustelid abundance and survival was not influenced as strongly by territory acquisition as was the relationship between lynx abundance and survival.

Survival of juveniles without a territory was higher in colder winters, with the opposite being true for juveniles holding a territory by autumn. We predicted that cold winters would lead to higher over winter mortality of territory owners, and we expected this to be magnified for non-territory owners, not reversed. There are some situations in which colder winters lead to higher survival, such as hibernating species (bats *Chalinolobus tuberculatus*; Pryde, O'Donnell, & Barker, 2005; jumping mice *Zapus hudsonicus preblei*; Schorr, Lukacs, & Florent, 2009) where this is thought to be due to less frequent arousal from hibernation (Humphries, Thomas, & Speakman, 2002). Red squirrels are non-hibernating, so this mechanism cannot explain why non-territory owners would benefit from colder winters.

We can suggest two alternative (but not mutually exclusive) explanations for why juveniles that acquired a territory late would have higher survival over winter. First, in colder years the incidence of nest-sharing among non-territory owners might be higher. Nest sharing, typically between kin, occurs in 19 % of female territory owners in this system, and is more common in colder winters (Williams et al., 2013). Juveniles without territories in autumn may be more likely to share nests with fellow non-territory owners, and this may improve their survival relative to juveniles with territories in autumn. Second, higher mortality of territory owners in colder winters creates vacancies, which may allow juveniles without territories by autumn a greater opportunity to claim a territory with plentiful cached food, enhancing survival (Dunham, Warner, & Lawson, 1995). This would give them relatively improved survival compared to warmer years where few territory owners would die. Which, if either, of these mechanisms accounts for the differential effect of winter temperatures remains to be tested.

We found both lynx and mustelid abundances were negatively associated with juvenile over winter survival. Previous work found that predation does not exert a strong influence on red squirrel populations in the boreal forest (Boonstra, Boutin, et al., 2001). However, in this study, the effects of annual lynx and mustelid abundance on juvenile survival (−0.68 and −0.32 for those without territories by autumn) were comparable in strength to the effect of cone availability (0.38), which is the primary driver of red squirrel population dynamics (LaMontagne et al., 2013). The relatively strong effects of predators on over winter survival in this study might appear contradictory to previous findings, but two distinctions can be made. First, overall population size and individual probability of survival are not directly comparable. While red squirrel population size may be dictated by the availability of food and territories, predation could still affect *which* individuals survive (“compensatory predation”; Errington, 1946). Second, this study was concerned with over winter survival of only juveniles, and predator population size had the strongest effect on the 39 % of juveniles that did not have territories by autumn. The probability of survival of these juveniles is already low, so variation in survival in this subset is not likely to have a large impact on the total population size.

We predicted that the effects of lynx and mustelid population sizes on juvenile survival would be mediated by the availability of their alternate prey. We did not find a significant interaction of either predator-prey pairing on red squirrel survival. One potential explanation for this could be that predator populations closely track their prey. For example, there were few years in our dataset with high predator and low prey densities with which to evaluate these interactions. Additionally, we grouped three mustelid species together, and they may respond differently to vole abundance. Furthermore, although lynx switch from hares to red squirrels when the former are rare (O’Donoghue, Boutin, Krebs, Zuleta, et al., 1998), lynx and mustelids may predate on juvenile red squirrels opportunistically if juveniles are more susceptible to predation regardless of alternate prey availability. Juveniles without territories by autumn could be more susceptible to this, if it occurs.

We did not anticipate that the population sizes of voles and hares would have significant effects themselves on over winter survival of juvenile red squirrels. High hare abundance was associated with increased juvenile survival, while years with high vole abundances had lower juvenile survival. Red squirrels will opportunistically predate on snowshoe hare leverets in the spring and summer (O’Donoghue, 1994), but this additional food source should not have a strong effect over winter. Voles are not in strong competition with juveniles for resources, given red squirrels access arboreal food sources unavailable to voles, and red-backed voles are broad omnivores, feeding on vegetation, fungi, and arthropods (Boonstra, Krebs, Gilbert, & Schweiger, 2001). These species’ population densities may covary with another factor that influences juvenile survival not included in our analysis, but what this factor might be remains unclear.

In our survival model, juveniles with higher growth rates were more likely to survive to spring, but birth date had no effect. Previous work in this population has observed strong selection on both birth date and growth rate in annual survival of juveniles (Dantzer et al., 2013; Fisher et al., 2017; McAdam & Boutin, 2003; Williams, Lane, Humphries, McAdam, & Boutin, 2014). In preliminary models not including territory ownership, there was a significant effect of birth date on over winter survival. Once accounting for territory ownership, birth date stopped being important. This implies that early born juveniles are likely to acquire a territory sooner, but there are no further benefits of birth date for survival over winter. Both earlier birth dates and higher growth rates are thought to be beneficial in territory acquisition, but there was still an effect of growth rate on over winter survival after accounting for territory ownership. Furthermore, larger juveniles in the autumn are more likely to survive to spring (Larivée et al. 2010). Among juveniles for which we have body mass measurements in August (*n* = 757), juveniles with higher relative growth rates were larger (β = 7.95 ± 1.61, *t* = 4.93, *P* < 0.001), but earlier birth dates also had a significant effect on body mass in August (β = −8.89 ± 1.25, *t* = −7.11, *P* < 0.001) so this does not explain why growth rate provides further benefits over winter but birth date does not. Presumably, growth rate may be associated with other life history and behavioural traits (Biro & Stamps, 2008; Réale et al., 2010; Stamps, 2007) that could affect over winter survival.

### Conclusions

We have identified how the timing of a life history event – territory acquisition – influences juvenile survival, and how it mediates biotic and abiotic factors that influence survival. This gives us insight into how one trait can affect the opportunity for selection on others, and therefore the routes through which organisms can arrive at “fit” phenotypes. We encourage more researchers to study key life stages such as the juvenile period, when survival can be highly variable and so the opportunity for selection high, to better understand how traits are selected in populations. As this study was primarily concerned with over winter dynamics, investigations of juveniles during territory prospection and before settlement, and which traits or conditions are associated with territorial acquisition, would be informative in further explaining the mechanisms behind some of the patterns we observed.

## Acknowledgements

We acknowledge that the lands on which we have conducted our research for the past 30 years are within the traditional territory of the Champagne and Aishihik First Nations. We thank Agnes MacDonald for providing access to her trapline. We thank the many people that have been involved in data collection over the years within the Kluane Red Squirrel Project, and many thanks to Charles Krebs and the Kluane Boreal Forest Ecosystem Project and Community Ecological Monitoring Program for making their data freely available to the public. The Natural Sciences and Engineering Council of Canada, the National Science Foundation, and the Northern Scientific Training Program provided research support. We have no conflicts of interest. This is paper number XX of the Kluane Red Squirrel Project.

## Author contributions

JH and DF conceived the ideas and conducted the analyses; JH, DF, and ARM led the writing of the manuscript; SB, BD, JL, and AGM managed long term data collection and revised initial drafts and analyses. All authors contributed critically to the drafts and gave final approval for publication.

## Data accessibility

Data used to evaluate juvenile over winter survival, along with code to recreate analyses and figures, will be made available on Dryad upon publication.

## Literature Cited

Bates, D., Maechler, M., Bolker, B., & Walker, S. (2015). Fitting linear mixed-effects models using lme4. Journal of Statistical Software, 67:1–8. doi:10.18637/jss.v067.i01

Berteaux, D., & Boutin, S. (2000). Breeding dispersal in female North American red squirrels. Ecology, 81(5), 1311–1326. Retrieved from http://doi.wiley.com/10.1890/0012-9658(2000)081[1311:BDIFNA]2.0.CO;2

Biro, P. A., & Stamps, J. A. (2008). Are animal personality traits linked to life-history productivity? Trends in Ecology & Evolution, 23(7), 361–368. doi:10.1016/j.tree.2008.04.003

Boonstra, R., Boutin, S., Byrom, A., Karels, T. I. M., Hubbs, A., Stuart-smith, K., … Antpoehler, S. (2001). The Role of Red Squirrels and Arctic Ground Squirrels. Ecosystem Dynamics of the Boreal Forest: The Kluane Project, 179–215.

Boonstra, R., & Krebs, C. J. (2006). Population limitation of the northern red-backed vole in the boreal forests of northern Canada. Journal of Animal Ecology, 75(6), 1269–1284. doi:10.1111/j.1365-2656.2006.01149.x

Boonstra, R., Krebs, C. J., Gilbert, B., & Schweiger, S. (2001). Voles and Mice. In C. J. Krebs, S. Boutin, & R. Boonstra (Eds.), Ecosystem Dynamics of the Boreal Forest: The Kluane Project (pp. 215–239). New York: Oxford University Press.

Boutin, S., Krebs, C. J., Boonstra, R., Dale, M. R. T., Hannon, S. J., Martin, K., … Schweiger, S. (1995). Population changes of the vertebrate community during a snowshoe hare cycle in Canada’s boreal forest. Oikos, 74(1), 69. doi:10.2307/3545676

Boutin, S., & Schweiger, S. (1988). Manipulation of intruder pressure in red squirrels (Tamiasciurus hudsonicus): effects on territory size and acquisition. Canadian Journal of Zoology, 66(10), 2270–2274. doi:10.1139/z88-337

Carpenter, F. L. (1987). Food abundance and territoriality: To defend or not to defend? Integrative and Comparative Biology, 27(2), 387–399. doi:10.1093/icb/27.2.387

Clarke, M. F., Burke, K., Lair, H., Pocklington, R., Clarke, M. F., Burke, K., … Robert L. (1993). Familiarity affects escape behaviour of the Eastern chipmunk, Tamias striatus. Oikos, 66(3), 533–537.

Cooper, E. B., Taylor, R. W., Kelley, A. D., Martinig, A. R., Boutin, S., Humphries, M. M., … McAdam A. G. (2017). Personality is correlated with natal dispersal in North American red squirrels (Tamiasciurus hudsonicus). Behaviour. doi:10.1163/1568539X-00003450

Cowlishaw, G. (1997). Refuge use and predation risk in a desert baboon population. Animal Behaviour, 54(2), 241–253. doi:10.1006/anbe.1996.0466

Dantzer, B., Newman, A. E. M., Boonstra, R., Palme, R., Boutin, S., Humphries, M. M., & McAdam, A. G. (2013). Density triggers maternal hormones that increase adaptive offspring growth in a wild mammal. Science, 340(6137), 1215–1217. doi:10.1126/science.1235765

Descamps, S., Boutin, S., Berteaux, D., & Gaillard, J.-M. (2008). Age-specific variation in survival, reproductive success and offspring quality in red squirrels: evidence of senescence. Oikos, 117(9), 1406–1416. doi:10.1111/j.0030-1299.2008.16545.x

Dunham, M. L., Warner, R. R., & Lawson, J. W. (1995). The dynamics of territory acquisition: a model of two coexisting strategies. Theoretical Population Biology, 47, 347–364.

Errington, P. L. (1946). Predation and Vertebrate Populations. The Quarterly Review of Biology, 21(2), 144–177. doi:10.1086/395220

Everett, R. A., & Ruiz, G. M. (1993). Coarse woody debris as a refuge from predation in aquatic communities. An experimental test. Oecologia, 93(4), 475–486.

Ferguson, G. W., & Fox, S. F. (1984). Annual variation of survival advantage of large juvenile side-blotched lizards, Uta stansburiana: its causes and evolutionary significance. Evolution, 38(2), 342–349.

Fisher, D. N., Boutin, S., Dantzer, B., Humphries, M. M., Lane, J. E., & McAdam, A. G. (2017). Multilevel and sex-specific selection on competitive traits in North American red squirrels. Evolution, 71(7), 1841–1854. doi:10.1111/evo.13270

Fisher, D. N., Haines, J. A., Boutin, S., Dantzer, B., Lane, J. E., Coltman, D. W., & McAdam, A. G. (2019). Indirect effects on fitness between individuals that have never met via an extended phenotype. Ecology Letters. doi:10.1111/ele.13230

Fletcher, Q. E., Boutin, S., Lane, J. E., LaMontagne, J. M., McAdam, A. G., Krebs, C. J., & Humphries, M. M. (2010). The functional response of a hoarding seed predator to mast seeding. Ecology, 91(9), 2673–2683. doi:10.1890/09-1816.1

Fletcher, Q. E., Landry-Cuerrier, M., Boutin, S., McAdam, A. G., Speakman, J. R., & Humphries, M. M. (2013). Reproductive timing and reliance on hoarded capital resources by lactating red squirrels. Oecologia, 173(4), 1203–1215. doi:10.1007/s00442-013-2699-3

Fuller, W. A., Stebbins, L. L., & Dyke, G. R. (1969). Overwintering of small mammals near Great Slave Lake Northern Canada. Arctic, 22(1), 34–55. doi:10.2307/40507757

Gaillard, J.-M., Festa-Bianchet, M., Yoccoz, N. G., Loison, A., & Toïgo, C. (2000). Temporal Variation in Fitness Components and Population Dynamics of Large Herbivores. Annual Review of Ecology and Systematics, 31(1), 367–393. doi:10.1146/annurev.ecolsys.31.1.367

Gaillard, J. M., Festa-Bianchet, M., & Yoccoz, N. G. (1998). Population dynamics of large herbivores: Variable recruitment with constant adult survival. Trends in Ecology and Evolution, 13(2), 58–63. doi:10.1016/S0169-5347(97)01237-8

Garrett, M. G., & Franklin, W. L. (1988). Behavioral Ecology of Dispersal in the Black-Tailed Prairie Dog. Journal of Mammalogy, 69(2), 236–250. doi:10.2307/1381375

Goheen, J. R., & Swihart, R. K. (2005). Resource selection and predation of North American red squirrels in deciduous forest fragments. Journal of Mammalogy, 86(1), 22–28. doi:10.1644/1545-1542(2005)086<0022:rsapon>2.0.co;2

Greenwood, P. J., & Harvey, P. H. (1982). The natal and breeding dispersal of birds. Annual Review of Ecology and Systematics, 13, 1–21.

Haines, J. A., Coltman, D. W., Dantzer, B., Gorrell, J. C., Humphries, M. M., Lane, J. E., … Boutin S. (2018). Sexually selected infanticide by male red squirrels in advance of a mast year. Ecology. doi:10.1002/ecy.2158

Humphries, M. M., & Boutin, S. (2000). The Determinants of Optimal Litter Size in Free-Ranging Red Squirrels. Ecology, 81(10), 2867–2877. Retrieved from http://www.jstor.org/stable/177347?origin=crossref

Humphries, M. M., Thomas, D. W., & Speakman, J. R. (2002). Climate-mediated energetic constraints on the distribution of hibernating mammals. Nature, 418(6895), 313–316. doi:10.1038/nature00828

Jackson, D. M., Trayhurn, P., & Speakman, J. R. (2001). Associations between energetics and over-winter survival in the short-tailed field vole Microtus agrestis. Journal of Animal Ecology, 70(4), 633–640. doi:10.1046/j.1365-2656.2001.00518.x

Kerr, T. D., Boutin, S., LaMontagne, J. M., McAdam, A. G., & Humphries, M. M. (2007). Persistent maternal effects on juvenile survival in North American red squirrels. Biology Letters, 3(3), 289–291. doi:10.1098/rsbl.2006.0615

Kerr, T., & Descamps, S. (2008). Why do North American red squirrel, *Tamiasciurus hudsonicus*, mothers relocate their young? A predation-based hypothesis. Canadian Field-Naturalist, 122(1), 65–66. doi:10.1258/la.2010.010166

Krebs, C. J. (2001). General Introduction. In C. J. Krebs, S. Boutin, & R. Boonstra (Eds.), Ecosystem Dynamics of the Boreal Forest: The Kluane Project (pp. 3–8). New York: Oxford University Press. doi:10.1016/S0006-3495(64)86921-6

Lair, H. (1990). The calls of the red squirrel — a contextual analysis of function. Behaviour, 115(3), 254–282. doi:10.1163/156853990X00608

LaMontagne, J. M., Peters, S., & Boutin, S. (2005). A visual index for estimating cone production for individual white spruce trees. Canadian Journal of Forest Research, 35(12), 3020–3026. doi:10.1139/x05-210

LaMontagne, J. M., Williams, C. T., Donald, J. L., Humphries, M. M., McAdam, A. G., & Boutin, S. (2013). Linking intraspecific variation in territory size, cone supply, and survival of North American red squirrels. Journal of Mammalogy, 94(5), 1048–1058. doi:10.1644/12-MAMM-A-245.1

Lane, J. E., McAdam, A. G., Charmantier, A., Humphries, M. M., Coltman, D. W., Fletcher, Q., … Boutin S. (2015). Post-weaning parental care increases fitness but is not heritable in North American red squirrels. Journal of Evolutionary Biology, 28(6), 1203–12. doi:10.1111/jeb.12633

Larivée, M. L., Boutin, S., Speakman, J. R., McAdam, A. G., & Humphries, M. M. (2010). Associations between over-winter survival and resting metabolic rate in juvenile North American red squirrels. Functional Ecology, 24(3), 597–607. doi:10.1111/j.1365-2435.2009.01680.x

Larsen, K. W., & Boutin, S. (1994). Movements, survival, and settlement of red squirrel (*Tamiasciurus hudsonicus*) offspring. Ecology, 75(1), 214–223. doi:10.2307/1939395

McAdam, A. G., & Boutin, S. (2003). Variation in viability selection among cohorts of juvenile red squirrels (Tamiasciurus hudsonicus). Evolution; International Journal of Organic Evolution, 57(7), 1689–1697. doi:10.1111/j.0014-3820.2003.tb00374.x

McAdam, A. G., Boutin, S., Sykes, A. K., & Humphries, M. M. (2007). Life histories of female red squirrels and their contributions to population growth and lifetime fitness. Ecoscience, 14(3), 362. doi:10.2980/1195-6860(2007)14[362:LHOFRS]2.0.CO;2

Merritt, J. F. (1986). Winter Survival Adaptations of the Short-Tailed Shrew (Blarina brevicauda) in an Appalachian Montane Forest. Journal of Mammalogy, 67(3), 450–464. doi:10.2307/1381276

Metzgar, L. H. (1967). An experimental comparison of screech owl predation on resident and transient white-footed mice (Peromyscus leucopus). Journal of Mammalogy, 48(3), 387–391. doi:10.2307/1377771

Morrison, S. F., Pelchat, G., Donahue, A., & Hik, D. S. (2009). Influence of food hoarding behavior on the over-winter survival of pikas in strongly seasonal environments. Oecologia, 159(1), 107–116. doi:10.1007/s00442-008-1197-5

O’Connor, C. M., Norris, D. R., Crossin, G. T., & Cooke, S. J. (2014). Biological carryover effects: linking common concepts and mechanisms in ecology and evolution. Ecosphere, 5(3), 28.

O’Donoghue, M. (1994). Early survival of juvenile snowshoe hares. Ecology, 75(6), 1582–1592.

O’Donoghue, M., Boutin, S., Hofer, E. J., & Boonstra, R. (2001). Other mammalian predators. In C. J. Krebs, S. Boutin, & R. Boonstra (Eds.), Ecosystem Dynamics of the Boreal Forest: The Kluane Project (pp. 325–336). New York: Oxford University Press.

O’Donoghue, M., Boutin, S., Krebs, C. J., Murray, D. L., & Hofer, E. J. (1998). Behavioural responses of coyotes and lynx to the snowshoe hare cycle. Oikos, 82, 169–183.

O’Donoghue, M., Boutin, S., Krebs, C. J., Zuleta, G., Dennis, L., Donoghue, M. O., … Hofer E. J. (1998). Functional responses of coyotes and lynx to the snowshoe hare cycle. Ecology, 79(4), 1193–1208. doi:10.1890/0012-9658(1998)079[1193:FROCAL]2.0.CO;2

Oli, M. K., & Dobson, F. S. (2003). The Relative Importance of Life History Variables to Population Growth Rate in Mammals: Cole’s Prediction Revisited. The American Naturalist, 161(3), 422–440. doi:10.1086/367591

Pryde, M. A., O’Donnell, C. F. J., & Barker, R. J. (2005). Factors influencing survival and long-term population viability of New Zealand long-tailed bats (Chalinolobus tuberculatus): Implications for conservation. Biological Conservation, 126(2), 175–185. doi:10.1016/J.BIOCON.2005.05.006

Réale, D., Garant, D., Humphries, M. M., Bergeron, P., Careau, V., & Montiglio, P.-O. (2010). Personality and the emergence of the pace-of-life syndrome concept at the population level. Philosophical Transactions of the Royal Society of London. Series B, Biological Sciences, 365(1560), 4051–4063. doi:10.1098/rstb.2010.0208

Rödel, H. G., Bora, A., Kaetzke, P., Khaschei, M., Hutzelmeyer, H., & von Holst, D. (2004). Over-winter survival in subadult European rabbits: weather effects, density dependence, and the impact of individual characteristics. Oecologia, 140(4), 566–576. doi:10.1007/s00442-004-1616-1

Rödel, H. G., Zapka, M., Talke, S., Kornatz, T., Bruchner, B., & Hedler, C. (2015). Survival costs of fast exploration during juvenile life in a small mammal. Behavioral Ecology and Sociobiology, 69(2), 205–217. doi:10.1007/s00265-014-1833-5

Rodríguez, S., van Noordwijk, A. J., Álvarez, E., & Barba, E. (2016). A recipe for postfledging survival in great tits Parus major: be large and be early (but not too much). Ecology and Evolution, 6(13), 4458–4467. doi:10.1002/ece3.2192

Schielzeth, H. (2010). Simple means to improve the interpretability of regression coefficients. Methods in Ecology and Evolution, 1(2), 103–113. doi:10.1111/j.2041-210X.2010.00012.x

Schorr, R. A., Lukacs, P. M., & Florant, G. L. (2009). Body Mass and Winter Severity as Predictors of Overwinter Survival in Preble’s Meadow Jumping Mouse. Journal of Mammalogy, 90(1), 17–24. doi:10.1644/07-MAMM-A-392.1

Searcy, S. P., & Sponaugle, S. (2001). Selective mortality during the larval – juvenile transition in two coral reef fishes. Ecology, 82(9), 2452–2470. doi:10.1890/0012-9658(2001)082[2452:SMDTLJ]2.0.CO;2

Smith, C. C. (1968). The adaptive nature of social organization in the genus of tree squirrels Tamiasciurus. Ecological Monographs, 38(1), 31–64.

Stamps, J. A. (2007). Growth-mortality tradeoffs and “personality traits” in animals. Ecology Letters, 10(5), 355–363. doi:10.1111/j.1461-0248.2007.01034.x

Steele, M. A. (1998). Tamiasciurus hudsonicus. Mammalian Species, 586(586), 1–9. doi:10.1890/0012-9623(2004)85

Stuart-Smith, A. K., & Boutin, S. (1995). Predation on red squirrels during a snowshoe hare decline. Canadian Journal of Zoology, 73(4), 713–722. doi:10.1139/z95-083

Studd, E. K., Boutin, S., McAdam, A. G., Krebs, C. J., & Humphries, M. M. (2015). Predators, energetics and fitness drive neonatal reproductive failure in red squirrels. Journal of Animal Ecology, 84(1), 249–259. doi:10.1111/1365-2656.12279

van der Jeugd, H., & Larsson, K. (1998). Pre-breeding survival of barnacle geese Branta leucopsis in relation to fledgling characteristics. Journal of Animal Ecology, 67, 953–966.

Whitham, T. G. (1986). Costs and benefits of territoriality: behavioral and reproductive release by competing aphids. Ecology, 67(1), 139–147. doi:10.2307/1938512

Williams, C. T., Gorrell, J. C., Lane, J. E., McAdam, A. G., Humphries, M. M., & Boutin, S. (2013). Communal nesting in an ‘asocial’ mammal: social thermoregulation among spatially dispersed kin. Behavioral Ecology and Sociobiology, 67(5), 757–763. doi:10.1007/s00265-013-1499-4

Williams, C. T., Lane, J. E., Humphries, M. M., McAdam, A. G., & Boutin, S. (2014). Reproductive phenology of a food-hoarding mast-seed consumer: Resource- and density-dependent benefits of early breeding in red squirrels. Oecologia, 174(3), 777–788. doi:10.1007/s00442-013-2826-1

Wunder, B. A., Dobkin, D. S., & Gettinger, R. D. (1977). Shifts of thermogenesis in the prairie vole. Oecologia, 29, 11–26.

