## Supplementary materials - Table S1 for "Territory acquisition mediates the influence of predators and climate on juvenile red squirrel survival"

### Supporting Information

Table S1. Mixed effects binomial model of juvenile red squirrel over winter survival (*n* = 1305), testing whether birth date or nestling growth rate mediate the effects of predators and temperature over winter, including random effects of litter ID and year.

| Term | Estimate ± SE | *z* | *P* |
| --- | --- | --- | --- |
| Std. density | -0.67 ± 0.16 | -4.17 | < 0.001 |
| Std. cones | 0.39 ± 0.13 | 3.00 | 0.002 |
| Std. growth rate | 0.29 ± 0.11 | 2.61 | 0.009 |
| Std. birth date | 0.01 ± 0.09 | 0.10 | 0.924 |
| Grid (SU) | 0.22 ± 0.18 | 1.20 | 0.229 |
| Sex (male) | -0.53 ± 0.15 | -3.44 | < 0.001 |
| Autumn owner (yes) | 2.67 ± 0.21 | 12.60 | < 0.001 |
| Std. lynx | 0.17 ± 0.22 | 0.78 | 0.433 |
| Std. hares | 0.13 ± 0.21 | 0.61 | 0.542 |
| Std. mustelid | -0.21 ± 0.13 | -1.65 | 0.099 |
| Std. voles | -0.51 ± 0.14 | -3.53 | < 0.001 |
| Std. temperature | 0.46 ± 0.13 | 3.52 | < 0.001 |
| Std. lynx : Std. hares | 0.12 ± 0.10 | 1.22 | 0.223 |
| Std. mustelid : Std. voles | 0.12 ± 0.13 | 0.92 | 0.357 |
| Std. growth rate : Std. lynx | -0.02 ± 0.13 | -0.15 | 0/882 |
| Std. growth rate: Std. mustelid | -0.12 ± 0.09 | -1.31 | 0.191 |
| Std. growth rate : Std. temperature | -0.09 ± 0.11 | -0.81 | 0.420 |
| Std. birth date : Std. lynx | -0.05 ± 0.10 | -0.49 | 0.623 |
| Std. birth date: Std. mustelid | 0.04 ± 0.08 | 0.49 | 0.623 |
| Std. birth date : Std. temperature | -0.03 ± 0.08 | -0.31 | 0.759 |
| Random effects | Variance | | |
| Litter ID | 0.544 | | |
| Year | 0.080 | | |
